# Role of Differential Expression of miRNAs miR-125a, miR-200a and miR-199a in Metastatic Property of Ovarian Cancer Cell lines and Relationship between miR-199a and its Predicted Target Protein GSK3β

**DOI:** 10.1101/2020.04.11.037424

**Authors:** Divya RSJB Rana, Lin Xu, Jie Zhang

## Abstract

Ovarian cancer is a gynecological cancer of high mortality rate. Most of the ovarian cancer origin from the surface epithelium of the ovaries. Ovarian cancers gain metastatic and invasive properties from various biochemical events taking place during tumorogenesis. MicroRNAs (miRNA) are non-coding RNAs and they have an important function of inhibiting translation of specific mRNAs in the cytoplasm. Present study uses two ovarian cancer cell lines as model to study the differential expression of miR-125a, miR-200a, and miR-199a, and provides indirect proof for possible relationship between miR-199a and its predicted target gene Glycogen Synthase Kinase 3 β (GSK3β). As expected the expression of these three microRNAs are decreased in metastatic ovarian cancer cell line A2780 compared to epithelial ovarian cancer OVCAR3. Reciprocal expression pattern of miR-199a and its predicted target GSK3β was found in two different cell lines providing information on possible inhibitory role of miR-199a against GSK3β.

## Introduction

Ovarian cancer is a type of gynaecological tumor with a high lethality, 90% of which occur in epithelial tissue on the surface of ovary, and the rest occuring in stromal and germ cells. Ovarian surface epithelial tissue are different form normal epithelial tissue in that they have mesenchymal cell characteristics (1). Although the ovarian epithelial tissue is of the same embryonic origin as the fallopian tubes and endometrial epithelium, this tissue has characteristics of normal epithelial tissues and can express E-cadherin (epithelial cadherin) and other keratin proteins (1). Ovarian epithelial cells will show normal epithelial characteristics after transformation into tumor cells, such as the expression of E-cadherin, while other epithelial tumor lose E-cadherin expression during tumor development (2).

MicroRNAs (miRNA) are a class of non-coding RNAs that inhibit translation initiation or cause target mRNA degradation based on complementarity of its sequence to that the 3’UTR of target mRNA, and play an important role in biological phenomena such as cell proliferation, differentiation, and apoptosis (3). Various miRNAs may be over- or under-expressed in ovarian tumors. MiRNA family miR-200 (4) and family miR-125/miR-125/let-7 are differentially expressed (4–5). Recent studies have shown that changes in expression of some miRNAs can promote epithelial-mesenchymal transition (EMT) (6). The process of tumor metastasis and invasion are closely related to the biochemical process of EMT. Decreased expression of miR-125a (7) and miR-200 family (8) promote transformation of ovarian tumor epithelial cells to mesenchymal cells, thereby increasing the ability to metastasize and invade. miR-199a has also been implicated in ovarian cancers, and studies have shown thati it can either inhibit the expression discoidin domain receptor 1 (DDR1) thus promoting the EMT process (9) or inhibit the EMT process (10). Expression of miR-199a has been found to be increased by transcription factor Twist which is found to increase metastatic property of cancers (11).

In this study, two ovarian cancer epithelial cell lines, OVCAR3 and A2780, were used as the research object to explore the differences in expression of three miRNA reported to be linked to metastatic properties of ovarian cancers, miR-125a, miR-200a, and miR-199a, and to explore the possible relationship between miR-199a and its predicted target gene glycogen synthase kinase 3 beta (GSK3β). The ovarian cancer cell line A2780 cell line has epithelial tissue morphology but is generally used for metastatic and invasive cell models (12,13). Park et al. has reported that the OVCAR3 cell line as a cell line with epithelial tissue characteristics (14) with higher E-cadherin expression and lower Vimentin expression, a protein expressed in invasive cells. Activation of GSK3β leads to apoptosis and its inactivation helps cell survival. GSK3β can directly lead to proteasome-dependent degradation of the Snail transcription factor (15). In addition, GSK3β can also inhibit the transcription of Snail through NF-ҝB (16). Many studies have demonstrated that Snail transcription factor can promote the transformation of epithelial tumor cells from epithelial cells to mesenchymal cells.

MicroRNA target prediction databases (MicroCosm, TargetScan and PicTar) using various algorithms *in silico* to predict target genes for published human miRNAs were used to detect the correlation between various miRNA and mRNA expressed in ovarian cancers. The three databases predicted that miR-199a interacts with GSK3β to varying degrees. This study hypothesized that expression of miR-125a, miR-200a and miR-199a are decreased in metastatic ovarian cancer cell lines. In addition, the inhibitory potential of miR-199a against GSK3β was explored through expressional differences at RNA and protein levels level.

## Material and Methods

### A. Cell culture

Two ovarian cancer cell lines A2780 and OVCAR3 were purchased from Chinese Acadmey of Sciences Cell Bank and cultured in high glucose DMEM medium or RPMI medium (Hyclone, USA) with 10% fetal bovine serum (Tianjin Yanyang, China) and 100 U/ml penicillin and 100 µg/ml streptomycin at 37°C and 5% carbon dioxide condition incubator (Thormer, USA). For sub-culturing of the cells, cells were first washed with calcium-free phosphate buffered-saline (PBS), digested with 0.25% trypsin in 0.1% ethylene diamine tetraacetic acid (EDTA) for 1-6 minutes (depending upon cell type) and divided into 1:2 to 1:4 proportions.

### B. PCR Detection of mRNAs and miRNAs

#### 1. Total RNA extraction

Cells fed with media within 2 days were washed in pre-chilled PBS and lysed in TRIzol (Invirogen, USA) (1ml per 10cm^2^ cells) for 5 minutes. After lysis, the solution was transferred to a clean 1.5 ml Eppendorf tube and added with chloroform (0.2 ml for each 1 ml of lysed solution). The mixture was vortexed for 15 seconds, incubated at room temperature (RT) for 3 minutes and spun at 12,000xg for 15 minutes at 4°C. The supernatant was transferred to RNAse-free Eppendorf tube (treated with DEPC), added with isopropanol (0.5ml for each 1 ml of supernatant), incubated at RT for 10 minutes and respun at 12,000xg for 10 minutes. The supernatant was discarded and the precipitate was washed with 1 ml of 70% ethanol and spun at 7,500xg for 5 minutes at 4°C. The supernatant ethanol was discarded and precipitate dried for 5-15 minutes and finally resuspended in 10-20µl DEPC-treated water. The RNA concentration was measured by spectrophotometer (1 µl RNA and 79 µl DEPC treated water) and remaining RNA was stored at −80C freezer. RNA was considered pure if optical density ratio (A260/A280) was within 1.8 and 2.0. RNA concentration was measured as:

Concentration: 40xA260xdillution factor ng/µl

#### 2. Reverse transcription (mRNA and miRNA)

Reverse transcription was performed using Primescript II 1st strand cDNA synthesis kit (TaKaRa, Japan) according to manufacturer’s instructions. Mixture of approximately □ 1.5 µg of template total RNA, 1µl oligo dT primer (50 mM) and 1 µl dNTP mixed solution (10 mM each) were constituted to 10µl with RNase-free water. After incubating at 65°C for 5 minutes, the solution was quickly cooled on ice. Then, 4 µl of 5× Reverse Transcription Buffer, 0.5 µl of RNase Inhibitor (20 U), 1 µl of Reverse Transcriptase (200 U) and RNase-free water were added to make 20 µl reaction volume. After mild vortex, the solution was incubated at 42°C for 60 minutes, 95°C for 5 minutes, and 4°C for 5 minutes, finally the cDNA obtained was stored at −30°C. The reverse transcription of miRNA was performed with total RNA. The procedure was as described above, but several steps were changed according to Tang et al. (17). The reverse transcription primers contain a 36 nucleotides long sequence forming hair-pin/stem-loop in the 5’ region followed by a short (approx. 8 nucleotides) long 3’ sequence to specifically bind the 3’ region of specific miRNAs. Reverse transcription of mature miRNAs were carried with 1µl 5′-terminal stem-loop special reverse transcription primers (10µM) to reverse transcribe specific miRNA (Table I) and 1µl small RNAU6 PCR downstream primers (10µM) to reverse transcribe U6 small RNA (Table II). U6 small RNA was used as an internal reference for miRNA expression detection. The reverse transcription reaction conditions are as follows: 16 □ for 30 minutes; 60 cycles of 20 □ for 30 seconds and 42 □ for 30 seconds, then, 50 □ for 1 second, 95 □ for 5 seconds and 4 □ standby temperature. The obtained first-strand cDNA was stored at −80 ° C.

**Table 1:**
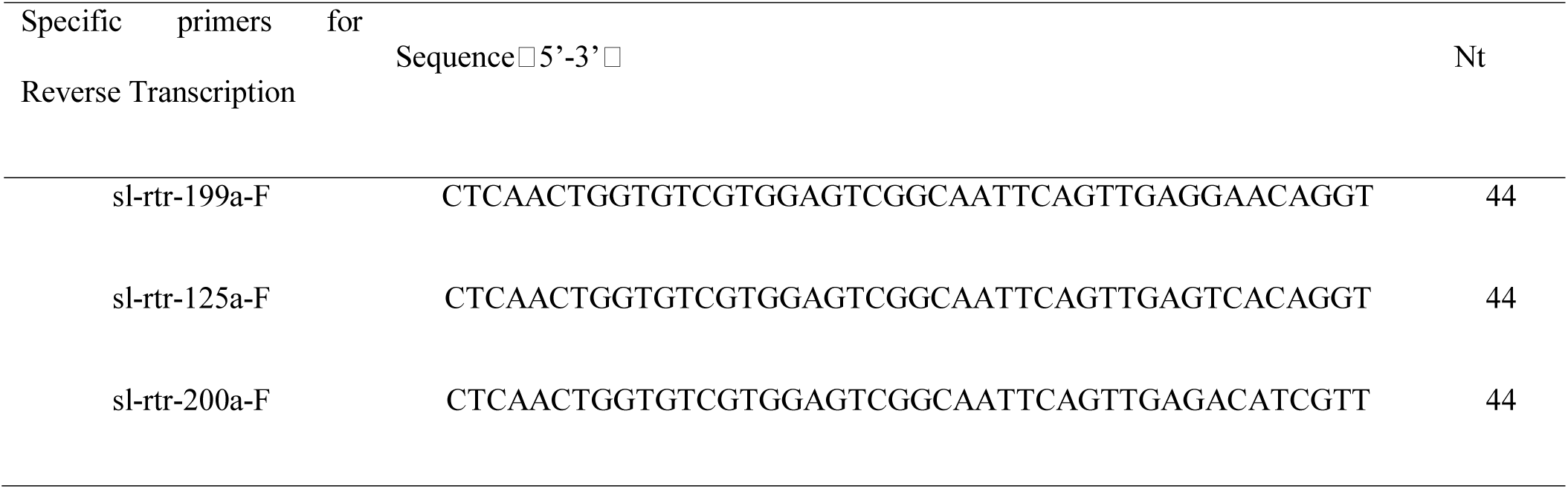
Specific Primers for Reverse Transcription of miRNAs

**Table 2:**
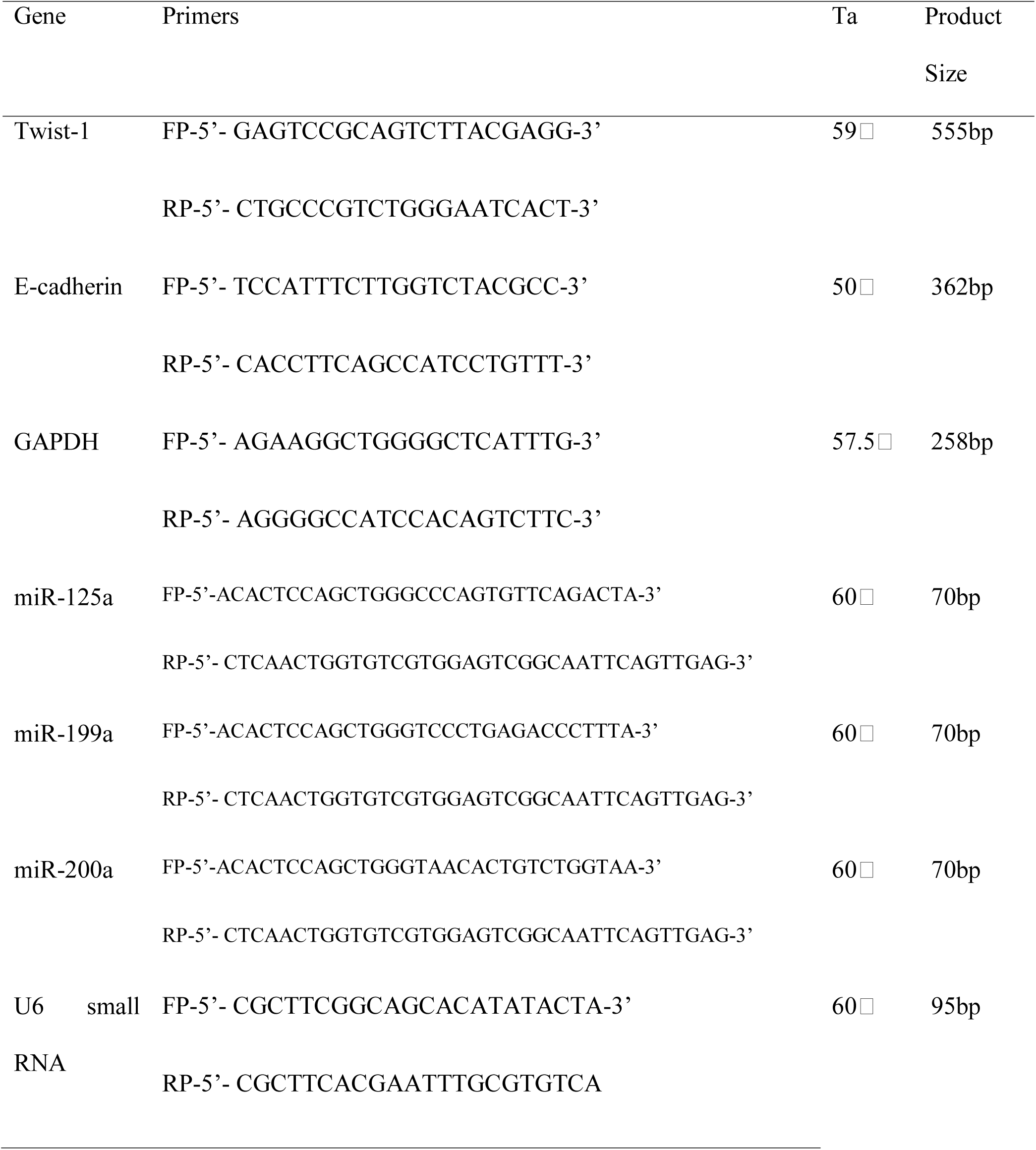
Primers for RT-conventional PCR for mRNAs and real time-PCR for miRNAs

#### 3. Reverse Transcription (RT)-PCR and real-time PCR

Reverse Transcription-conventional Polymerase Chain Reaction was used to detect mRNA levels of specific genes and real-time PCR to detect miRNA expression levels. The PCR primer sequences are shown in Table II. Premix Taq version 2.0 kit (TaKaRa, Japan) was used for PCR reaction. The reaction system was as follows: 25µl of 2× premix rTaq, 2/2 µl 10µM forward/reverse primers, 1µl cDNA (containing 50-75ng/µl RNA), and 20µl deionized water. The PCR reaction conditions were as follows: 94 □ for 5minutes, 25 to 30 cycles of 94 □ for 30seconds and 50-60 □ for 30 seconds and 72 □ for 5 minutes, finally stored at 4 L.

The Qiagen SYBR Green PCR kit (Qiagen, Germany) was used for real-time PCR. The reaction system was as follows: 12.5µl 2× Master Mix, 2.5/2.5µl forward/reverse miRNA-specific primers (10µM), 1µl cDNA template (50□75ng / µl) and water to 25µl. The 5′-end portion of all the forward primers is the same, the 3′-end portion is specific, and the reverse primer is the same/universal for all. The forward and reverse primers of the internal reference U6 PCR reaction system had 1 µl (10 µM) of each forward and reverse primers, and total volume was made up to 25 µl with water. ABI7500 real-time PCR instrument was used and the reaction conditions were as follows: 95 ° C for 15min, 40 cycles of two steps at 94 ° C for 15sec and 60 ° C for 1min.

#### 4. GSK3β protein level detection by Western Blot

A2780 and OVCAR3 cells were collected in Eppendorf tubes within 2 days of replacing growth media, washed twice with pre-chilled PBS, and centrifuged at 1000 rpm/min for 5 minutes.. The supernatant were discarded and 80 μL lysis buffer (20 mmol/ □ Tris / HCl pH 7.5, 50mmol/L NaCl, 0.1mmol/L Na3PO4, 25 mmol/L NaF, 2 mmol/L EDTA/EGTA, 1mmol/L DTT, 1 mg/L Leupeptidase/Aprotinin) were added for cells per culture bottle, placed in ice for 30 minutes, and sonicated for 20 seconds for 3-4 times. The solution was centrifuged at 12,000 rpm for 20 minutes and the supernatant was used as protein solution. Coomassie Brilliant Blue method was used to determine the protein content. 60 μg of total protein was loaded per well and separated by 10% SDS-PAGE gel electrophoresis and the proteins were transferred to a PVDF membrane. The membrane was cut according to molecular weight of GSK3β. The membrane was blocked with 5% skimmed milk powder at room temperature for 2 hours, incubated with primary antibody (1:500 of GSK3β, rabbit anti-human antibody, Beijing Biosynthesis, China) at 4°C overnight, and incubated again with the horseradish peroxidase-labeled secondary antibody (1:2000 of Goat anti-rabbit HRP labelled IgG, Santa Cruz, USA) at room temperature for 1 hour. After incubation, the membrane was eluted 3 times with TBS containing 5% Tween-20 for 10 min each. After drying, the PVDF membrane was developed with ECL. The gray value was scanned by gel imaging analysis system (ChemiBis 3.2, Israel).

#### 5. Statistical methods

The difference in miRNA or protein expression between the two cell lines was detected by t-test, with p ≤ 0.05 as the was statistically significant. The means and standard errors of mean shown in figures were calculated from triplicate experiments

## Results

### 1. Expressional analysis by reverse transcription convetional and real-time PCR E-cadherin and Twist1 mRNA expression levels detected by PCR

PCR results showed that E-cadherin expression was not detected in the ovarian cancer cell lines A2780, but detected in the OVCAR3 cell line (each PCR reaction was carried for 30 cycles with 50ng cDNA template) (Figure 1A). Twist1 mRNA expression was apparent in both A2780 and OVCAR3 cell lines (Figure 1B). GAPDH (25 cycles) could be amplified from the cDNAs from both of the cell lines

**Figure 1:**
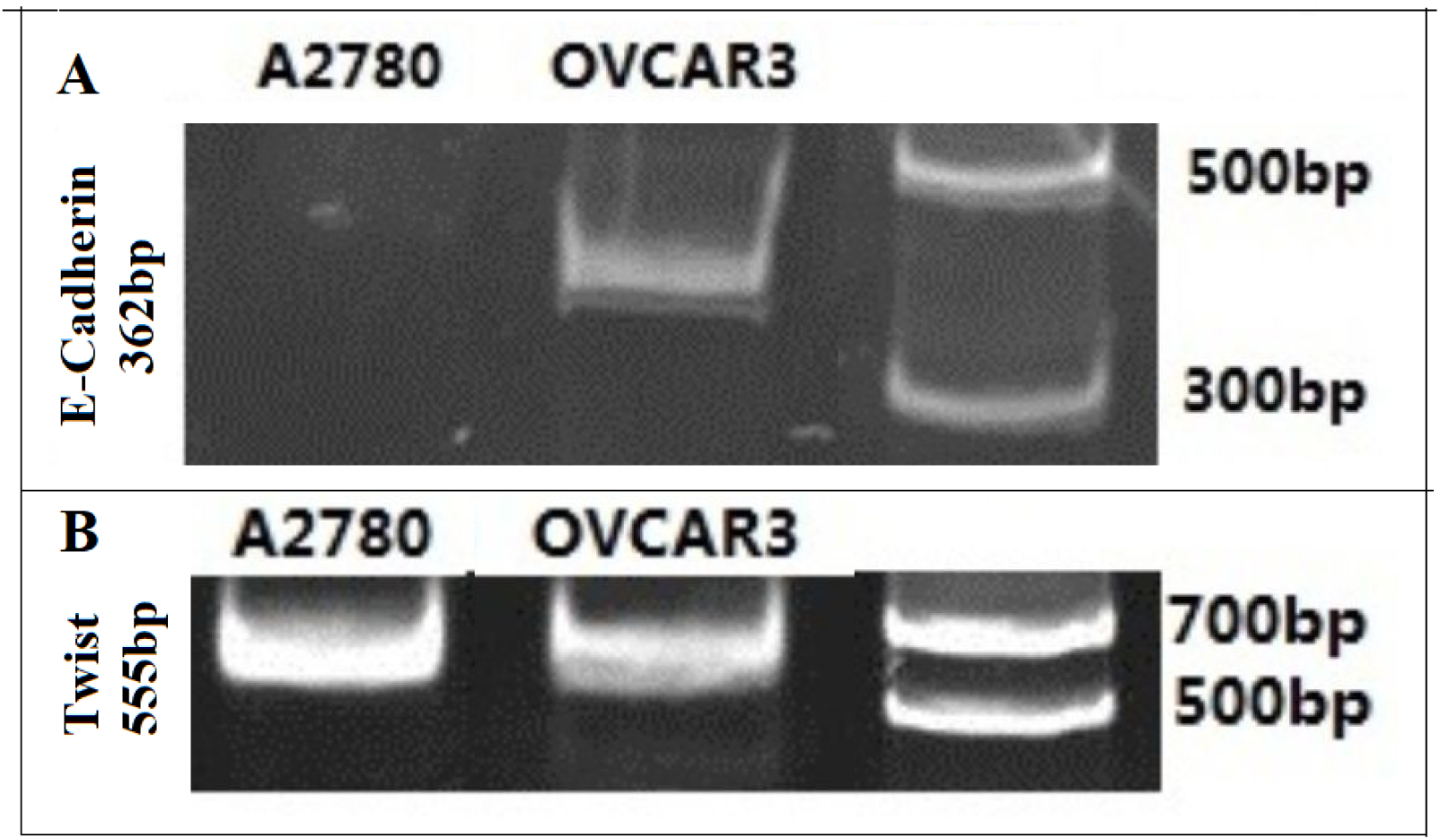
Expression of E-cadherin (362 bp) and Twist1 (555bp) PCR product in cell lines A2780 and OVCAR3

There were significant differences in the expression of different miRNAs in the two cell lines A2780 and OVCAR3 (Figure 2). miR-125a had the higher expression in OVCAR3 compared to A2780 (p <0.05). Similarly, miR-200a had the higher expression in OVCAR3 compared to A2780 (p <0.001) (Figure 2). There were statistically significant differences in miR-199a expression between OVCAR3 and A2780. miR-199a had higher expression in OVCAR3 than A2780 (p <0.0001).

**Figure 2:**
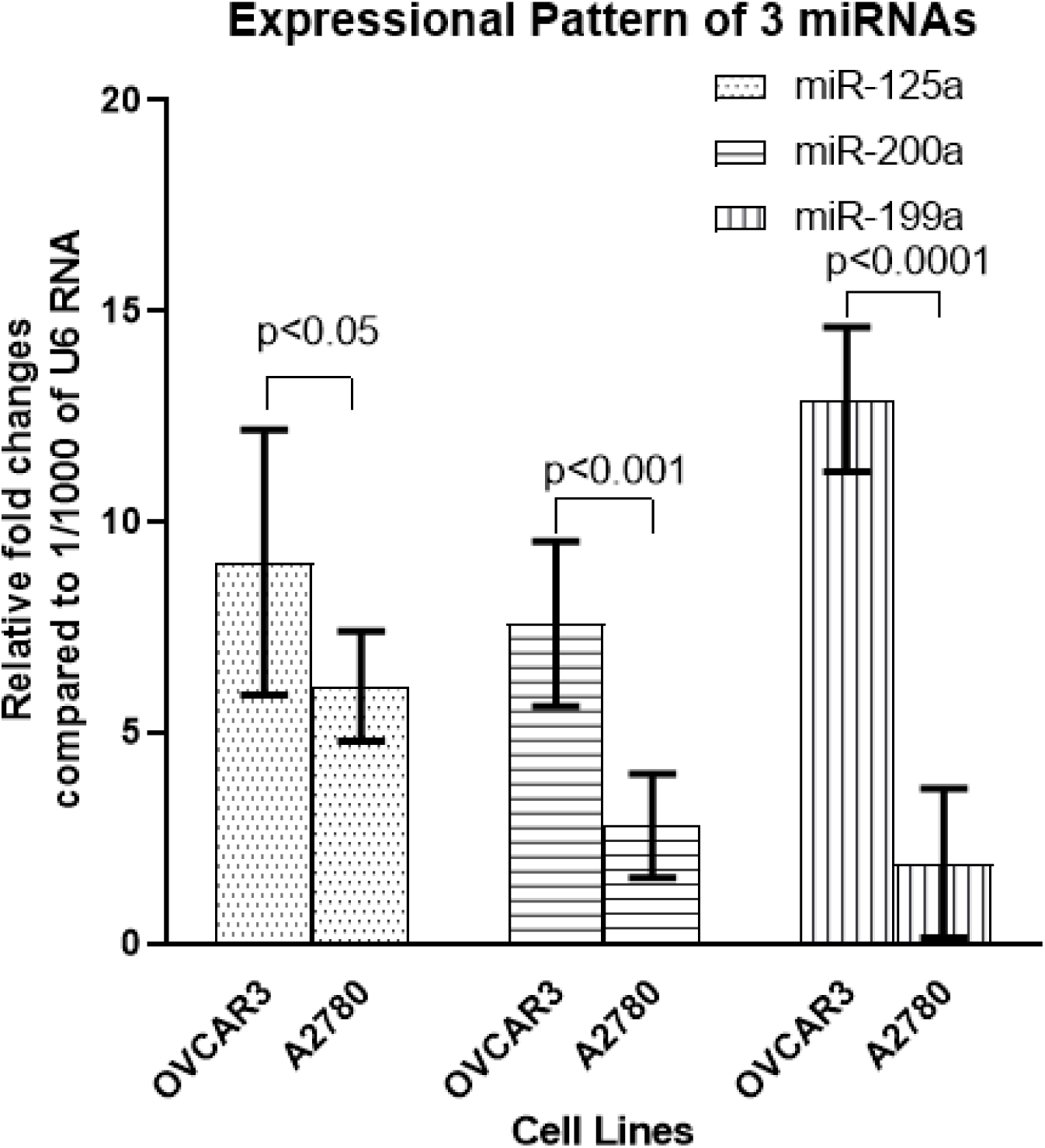
Expression of miR-125a, miR-200a and miR-199a in two ovarian cancer cell lines OVCAR3 and A2780. Bars show 95% confidence intervals. Experiments were done in triplicates.

### 2. Western blot detection of GSK3β protein levels

The difference in GSK3β protein between A2780 and OVCAR was statistically significant (p<0.05) (Figure 3). In this experiment, an equal volume and equal concentration of total protein-protein loading buffer mixture were loaded, and β-actin was not used as an internal reference.

**Figure 3:**
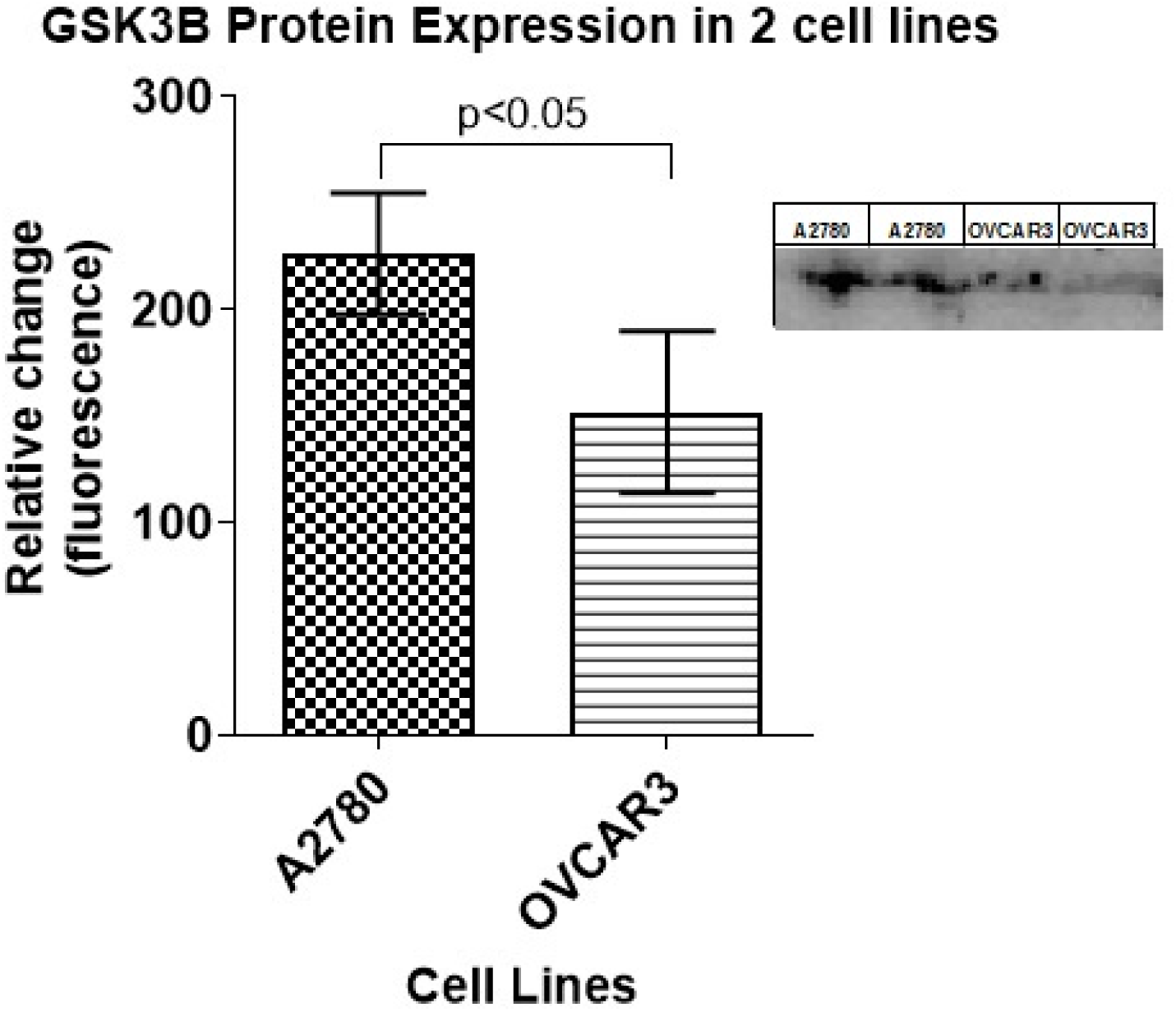
Expression of GSK3β by Western Blot in 2 ovarian cancer cell lines A2780 and OVCAR3. Bars show 95% confidence intervals. Experiments were done in duplicates.

## Discussion

E-cadherin was expressed in OVCAR3, and not in A2780. Our results are in line with previous studies which found that OVCAR3 are epithelial in nature and the A2780 two are metastatic (12,13 14). Twist1 is an important transcription factor that promotes EMT (18), and Yin and others study has shown that Twist1 protein can up-regulate the expression of miR-199a (11). This study shows that Twist1 is expressed in both A2780 and OVCAR3, but the expression of miR-199a in OVCAR3 is higher than in A2780. This phenomenon may be due to the presence of other proteins that affect miR-199a expressiion. Activation of Akt protein and the presence of STAT3 protein can inhibit miR-199a expression (19,20). Studies have shown that the high expression of miR-125a and miR-200 families is directly related to the epithelial characteristics of tumor cells. We found that both miR-125a and miR-200a had higher expression in epithelial cell line OVCAR3 compared to the metastatic cell line A2780, confirming their roles in metastasis of ovarian cancers. Some of the proteins that to promote EMT, like endothelin (21) and HIF1A (22), have been shown to be inhibited by miR-199a (32, now 23). In addition, EMT in pancreatic cancer was found to be a direct consequence of down-regulation of miR-199a, which increased the expression of DDR1 (9). Contrary, miR-199a also inhibits PIAS3 in cervical carcinoma to promote EMT (10). Thus, though decreased miR-199a may act as indicator for activation of EMT in tumor cells, in some contexts it can play opposite roles. Based on miRNA target databases we predicted that GSK3β is a target gene of miR-199a, and 3′-UTR of GSK3β has a target sequence of miR-199a. Western blot results showed that GSK3β was more highly expressed in ovarian cancer cell line A2780 than in OVCAR3. As there is reciprocal expression of miR-199a and its predicted target GSK3β, there could be a possible inhibitory relationship between these two genes. This is not a direct proof of their relationship but the relationship has been conclusively proven in other tissue systems (24, 25). However, activated GSK3β inhibits Snail activity by either leading to proteasome-dependent degradation of the Snail transcription factor and transfer Snail from the nucleus to the cytoplasm (15), or inhibition of transcription through NF-kB (16). As Snail expression is an indicator of metastatic capacity, GSK3β expression is expected to be down-regulated in cells with high metastatic properties.

The limitations of this study include use of primary antibodies against non-phophorylated GSK3β. As phosphorylation at different sites show different states of activation (26), use of antibodies against specific phosphorylated GSK3β can shed more light on the fate of GSK3β during high and low miR-199a expression. Similarly, further studies using recombinant reporter proteins with 3’UTR of GSK3β into 3’UTR of reporter protein can confirm inhibitory roles of miR-199a against GSK3β. Moreover studies in actual ovarian cancer tissues or ovarian cancer primary cell lines can help realize the actual relevance of these findings.

## Conclusion

As expected in the hypothesis, this study indicated that expression of miR-125a, miR-200a and miR-199a are decreased in metastatic ovarian cancer cell line A2780 compared to non-metastatic epithelial ovarian cancer cell line OVCAR3. The expression level of miR-199a was opposite to that of its predicted target gene GSK3β, which show possible inhibitory role of miR-199a over GSK3β in ovarian cancer cell lines.

## Acknowledgements

We are grateful to support provided by Dr. Wang Guoli, Dr. Wang Huaqin, Dr. Zong Zhihong, and Dr. An Liwen from Department of Biochemistry and Molecular Biology. We also thank Dr. Xu Yingqi from Department of Medical Genomics for her valuable inputs during the qPCR experiments. And finally, we would thank Dr. Han Feng and Hou Qiang for helping to do experiments.

